# Elucidating spatially-resolved changes in host signaling during *Plasmodium* liver-stage infection

**DOI:** 10.1101/2021.09.22.461346

**Authors:** Elizabeth K.K. Glennon, Tinotenda Tongogara, Veronica I. Primavera, Sophia M. Reeder, Ling Wei, Alexis Kaushansky

## Abstract

Upon transmission to the human host, *Plasmodium* sporozoites exit the skin, are taken up by the blood stream, and then travel to the liver where they infect and significantly modify a single hepatocyte. Low infection rates within the liver have made proteomic studies of infected hepatocytes challenging, particularly *in vivo*, and existing studies have been largely unable to consider how protein and phosphoprotein differences are altered at different spatial locations within the heterogeneous liver. Using digital spatial profiling, we characterized changes in host signaling during *Plasmodium yoelii* infection *in vivo* without disrupting the liver tissue, and measured variation between infected cells. Moreover, we measured alterations in protein expression around infected hepatocytes and identified a subset of CD163^+^ Kupffer cells that migrate towards infected cells during infection. These data offer the first insight into the heterogeneity of the infected hepatocyte in situ and provide insights into how the parasite may alter the local microenvironment to influence its survival and modulate immunity.

## Introduction

Upon introduction to the human host by the bite of an infectious mosquito, *Plasmodium* parasites migrate to the liver where they invade a hepatocyte and proceed to develop and replicate. Once parasites complete their development within the liver, thousands of individual merozoites egress from the host hepatocyte and migrate to the bloodstream where they invade erythrocytes and initiate the symptomatic blood stage of infection. The liver is often viewed as a uniform organ, however, factors such as oxygen and nutrient gradients lead to diverse cellular phenotypes and the formation of niches within the tissue [1, 2]. *Plasmodium* parasites traverse multiple hepatocytes before invading one [3–5] and preferentially invade both particular liver zones [6] and hepatocytes with specific phenotypes, such as high ploidy and particular surface receptor compositions [7, 8]. In addition to selecting particular hepatocytes for invasion, parasites modify the host cell throughout their development within the liver, including cell size [9], microtubule and organelle organization [10], and signaling cascades [11, 12].

The liver stage (LS) is a substantial bottleneck in *Plasmodium* infection, making it an attractive point for intervention. Attrition in parasite numbers occurs between injection at the skin, invasion of hepatocytes, and completion of development within the liver [13]. Heterogeneity among hepatocytes within and between individuals can exacerbate this attrition; the ability of hepatocytes to support *Plasmodium falciparum* and *Plasmodium vivax* infection varied extensively between individual human donors [14]. Experiments with genetically attenuated parasites demonstrated that parasites that successfully invade but die before completing LS infection can induce immunity and reduce susceptibility to subsequent infection [15].

Several global studies have been conducted to understand alterations that occur during LS infection which may be important for the maintenance of infection. Transcriptomic studies have demonstrated extensive changes in host gene expression that vary over the course of infection, however concordance among these studies has been low, perhaps due to differences in hepatocyte origin and time needed to sort infected cells [16, 17]. Protein and post-translational modification level screens have been conducted using reverse phase protein array (RPPA) in an *in vitro* model of *Plasmodium yoelii* infection [12], and to identify proteins that are differentially expressed between hepatocyte populations of differential susceptibility to LS infection [11, 18]. Several proteins and processes that were identified as altered in infected cells were also found to be important for LS infection (reviewed in [19]). A small RPPA screen of infected hepatocytes revealed a suppression of p53 levels which was then found to be critical for avoidance of host cell death and maintenance of LS infection *in vitro* and *in vivo* [12, 20]. RNA-sequencing of infected hepatocytes revealed upregulated expression of aquaporin-3 (AQP3). Follow-up studies identified AQP3 as essential for infection and implicated it in nutrient acquisition [21]. Some functional screens have also been done to identify host proteins that are important for LS infection including siRNA, CRISPR, and kinase screens [22–24]. However, large-scale proteomic studies of liver-stage infection have been hindered by low infection rates, on the order of 1% in vitro and 0.01% *in vivo* [25]. Additionally, *in vivo* studies traditionally involve sorting infected from uninfected cells and pooling all uninfected cells together, thereby losing the ability to link parasite biology to its microenvironment, or heterogeneity among uninfected cells to their spatial distribution within the liver and position relative to the infected cell.

## Results

To interrogate differences in host cell signaling in intact *Plasmodium*-infected liver tissue we chose to utilize Digital Spatial Profiling (DSP). DSP interrogates levels of total and phosphorylated proteins in defined regions of fixed tissue [26], thereby preserving spatial information and limiting sample processing that could induce artificial changes. To date, DSP has primarily been used to study heterogeneity within the tumor microenvironment which has been strongly linked to disease progression and treatment outcomes [27, 28]. Briefly, liver sections are scanned and regions of interest (ROIs) are selected based on staining with fluorescent markers. Slides are incubated with one of several panels of antibodies bound with a photocleavable linker to unique oligonucleotide barcode tags. UV light is shone on the defined ROIs, cleaving the oligo tags from bound antibodies which are collected and quantified using the nCounter system (Figure 1A). We infected BALB/c mice with 1 million *P. yoelii* sporozoites and allowed the infection to proceed for 44 hours. Liver sections (4 μm thick) from 7 infected and 8 uninfected mice were stained using an antibody directed against parasite protein *Py*HSP70 that was fused with Alexa Fluor 488. In parallel with fluorescent staining, liver sections were also incubated with a panel of 42 oligo-tagged antibodies against a variety of host proteins and/or post-translational modifications (Table S1). Slides were imaged and pooled ROIs that encompassed five infected cells (50 μm diameters) or corresponding uninfected regions were identified (Fig1A). Oligos were cleaved and collected from ROIs using the GeoMx Digital Spatial Profiler for quantification. We observed host proteins/post-translational modifications that were both up- and down-regulated in infected regions compared to uninfected mice (Fig 1B), some of which have been previously identified as altered upon, or important for, infection (Table S1)[11, 12, 23, 29–32]. Because we conducted multiple DSP runs with tissues on multiple slides, we wanted to compare the reproducibility of our results and identify the contribution of run-to-run variation to the observed variability. Liver sections from infected and uninfected mice were evenly distributed across slides and runs. Comparisons of the fold change in protein levels (infected over uninfected) between two runs (Fig S1A), as well as between two slides within a single run (Fig S1B), all gave strong linear correlations, suggesting the data from multiple experiments are comparable. Interestingly, the slope of the line between two separate runs (Fig S1A) was greater than one, suggesting that comparisons of the magnitude of change between runs should be interpreted with caution.

**Figure 1.**
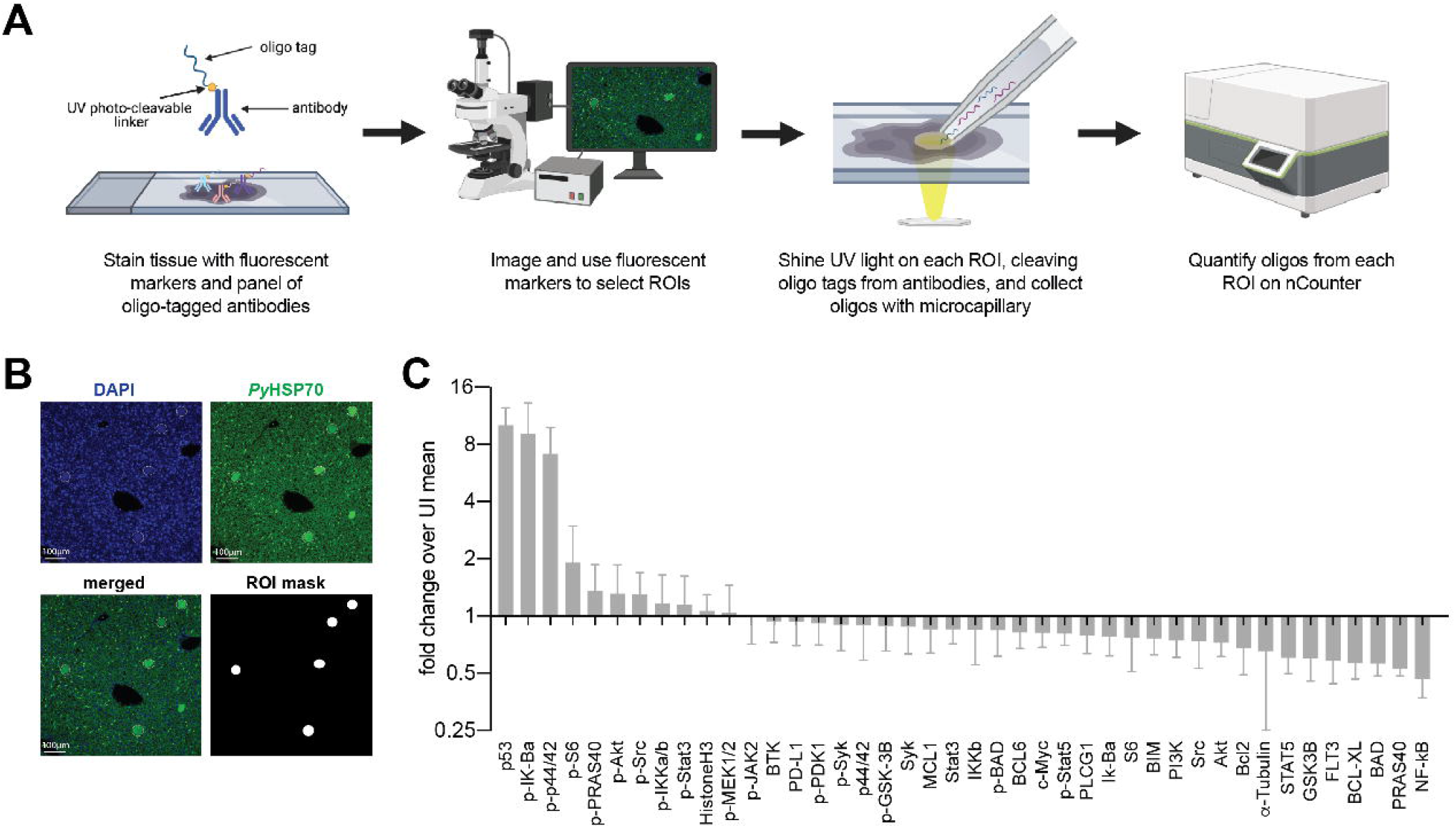
Digital Spatial Profiling facilitations evaluation of proteins and post-translational modifications in *Plasmodium yoelii* infected tissue. (A) Schematic of DSP methodology. (B) Representative images of fluorescent staining of liver sections from uninfected or *P. yoelii*-infected mice at 44hpi. Parasites are stained with *Py*HSP70 in green and DAPI is shown in blue. Data were pooled from regions of interest indicated by white circles. (C) Signal from *P. yoelii*-infected mice normalized to ROI area and to the average signal from uninfected mice. Error bars indicate standard deviation. n = 7-8 mice per group.

We next asked if we could reliably detect changes in host protein levels in single infected cells by DSP. We hypothesized that the enlarged size of infected hepatocytes at 44hpi (approximately 50μm in diameter) might allow us to reliably detect changes in host proteins at the single cell level. The same panel of antibodies (Table S1) was used to detect host protein levels in single infected-cell ROIs from a single infected mouse and in identical sized ROIs encompassing roughly 10 uninfected cells from a single uninfected mouse. Plotting the average of multiple infected single ROIs against the infected pooled ROI, for each antibody, gave a strong linear correlation (Fig S1C). The same was found for the comparable uninfected ROIs (Fig S1D), suggesting we have sufficient resolution to detect changes in host protein levels within single infected cells for these enlarged, infected cells. Upon examining the fold change between infected and uninfected single ROIs we observed a high degree of similarity between our pooled and single ROIs (Fig 2A). The same top four proteins were seen in both pooled and single ROIs, suggesting that the increase detected in the pooled ROIs was not due to a small population of high-expressing cells but may in fact be a feature of multiple infected hepatocytes. As an orthologous approach, we used immunofluorescent microscopy to evaluate several proteins that exhibited differential levels in infected and uninfected cells. Patterns of changes in protein levels between infected and uninfected ROIs were comparable as measured by DSP and by single fluorescent antibody staining (Fig 2B-C). We cannot currently rule out nonspecific binding to parasite proteins, however we hypothesize that the localization of several host proteins within the parasitophorous vacuole is due to uptake of host cell cytosol by the parasite, as has been described in blood stage parasites [33, 34].

**Figure 2.**
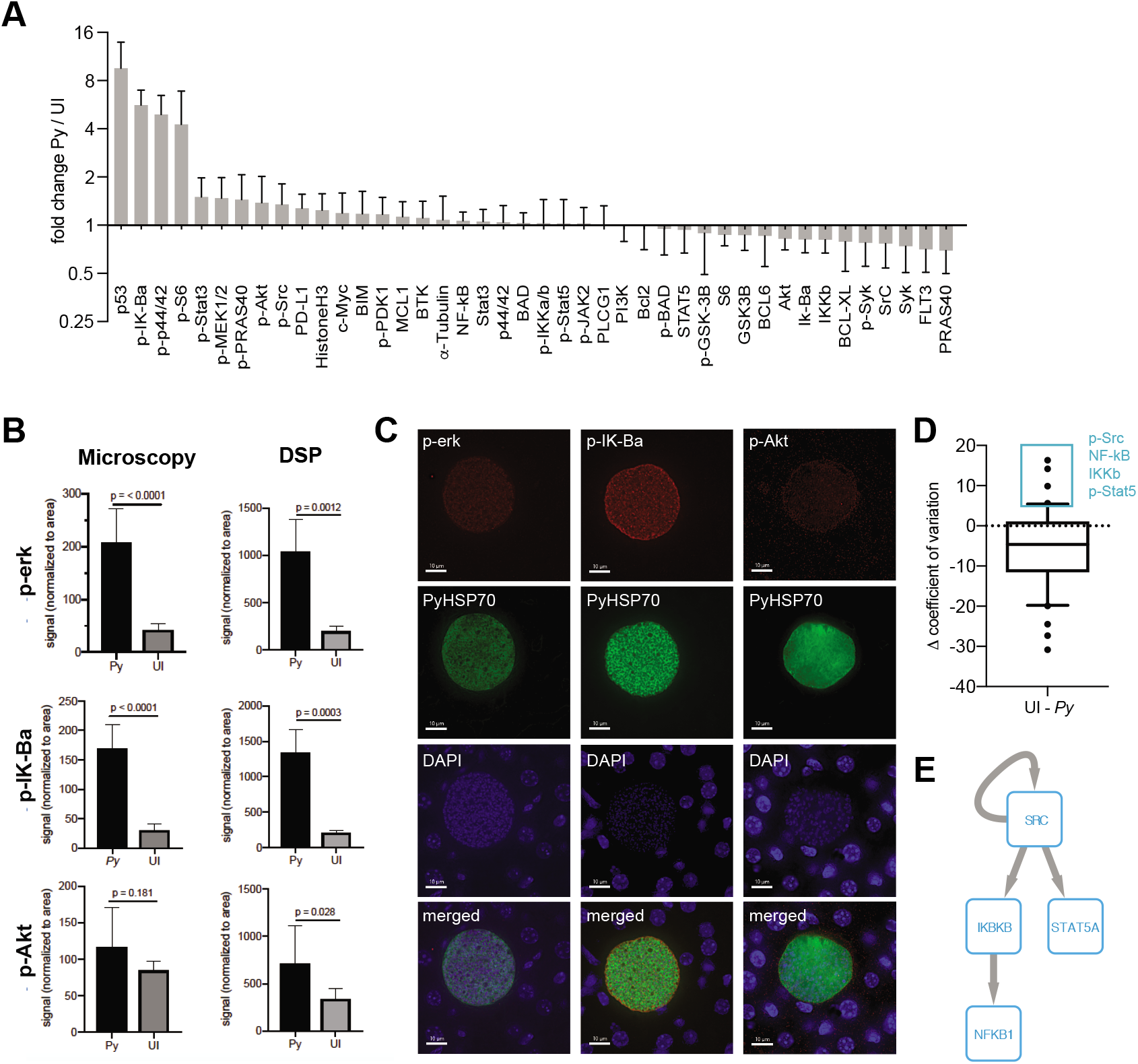
Single infected hepatocytes produce sufficient signal for detection by DSP. (A) Signal from 9 infected cells from a single *P. yoelii*-infected mouse normalized to region of interest (ROI) area and to the average signal from 6 uninfected cell regions from a single uninfected mouse. Error bars indicate standard deviation. (B) ROI-area-normalized signal from single infected and uninfected ROIs as measured by DSP (n = 6-9) and single antibody staining fluorescent microscopy (n = 6). (C) Representative images of infected cells were taken at a total magnification of 400x at 44hpi. (D) Difference in coefficient of variation between uninfected and single infected ROIs (9 infected cells) and 6 ROIs of identical size from a single mouse, for each antibody. Box plot encompasses 10-90^th^ percentile. Antibodies within the 90^th^ percentile which showed less variation in infected (*Py*) than in uninfected ROIs are boxed in blue. (E) Phosphosignaling network constructed from proteins with lower variation in infected ROIs. Arrows indicate direct phosphorylation events.

We used the ability to measure host protein levels in single infected cells to ask how host protein levels varied in single infected cells when compared to uninfected ROIs (Fig 2D, Table S2). We reasoned that proteins that exhibited substantially less variation between infected cells might represent features that are selected or tuned by the parasite to facilitate its survival and/or development. When examining the distribution of the difference in variation between infected and uninfected ROIs for our panel of antibodies, we observed a trend towards increased variation in infected ROIs (Fig 2D). This is likely explained by the masking of single cell variation within the uninfected ROI, which encompasses roughly 10 cells. Despite the skewedness of the distribution, several total and phosphorylated proteins (p-Src, NF-kB, IKKb, and p-Stat5) exhibited more variation in the uninfected ROI than in single infected cells (> 90^th^ percentile) (Fig 2D). To investigate whether or not these proteins might act as part of a connected network, we reconstructed a phosphosignaling network using a database of known kinase target sequences (Fig 2E). Network reconstruction revealed that proteins with lower variation in infected cells can directly interact with each other via phosphorylation, suggesting it could be a target of parasite selection and/or manipulation. Increased variation in infected cells could be due to host cell and/or parasite-intrinsic heterogeneity, or due to the influence of different local microenvironments within the liver.

We next investigated how areas surrounding infected cells varied with distance from the parasite. Using concentric ring ROIs matched with each LS parasite, we measured protein levels in infected cells, proximal uninfected cells (Ring1), and distal uninfected cells (Ring2) surrounding each parasite (Fig 3A). In addition to the original antibody panel, we included a second panel encompassing proteins expressed on various immune cells (Table S1). Protein levels in Ring1 and Ring2 were compared to those in their paired infected ROI and fell into clusters based on spatial patterns of relative expression (Fig 3B-C, Table S3). We were particularly interested in proteins with higher levels in Ring1 compared to Ring2 (Table S4) and theorized that these might be indicative of either (1) immune cell infiltration towards the infected hepatocyte, (2) selection of a cellular niche on a very fine scale, or (3) neighboring cells responding to signals emanating out from the infected cell. Of the proteins with significantly higher levels in Ring1 compared to Ring2, several immune cell surface markers, all of which have been described on macrophages (PD-L1, B7-H3, CD68, CD163) [35–37], were the most heavily upregulated (Table S4), leading us to investigate the distribution of macrophages around the parasite.

**Figure 3.**
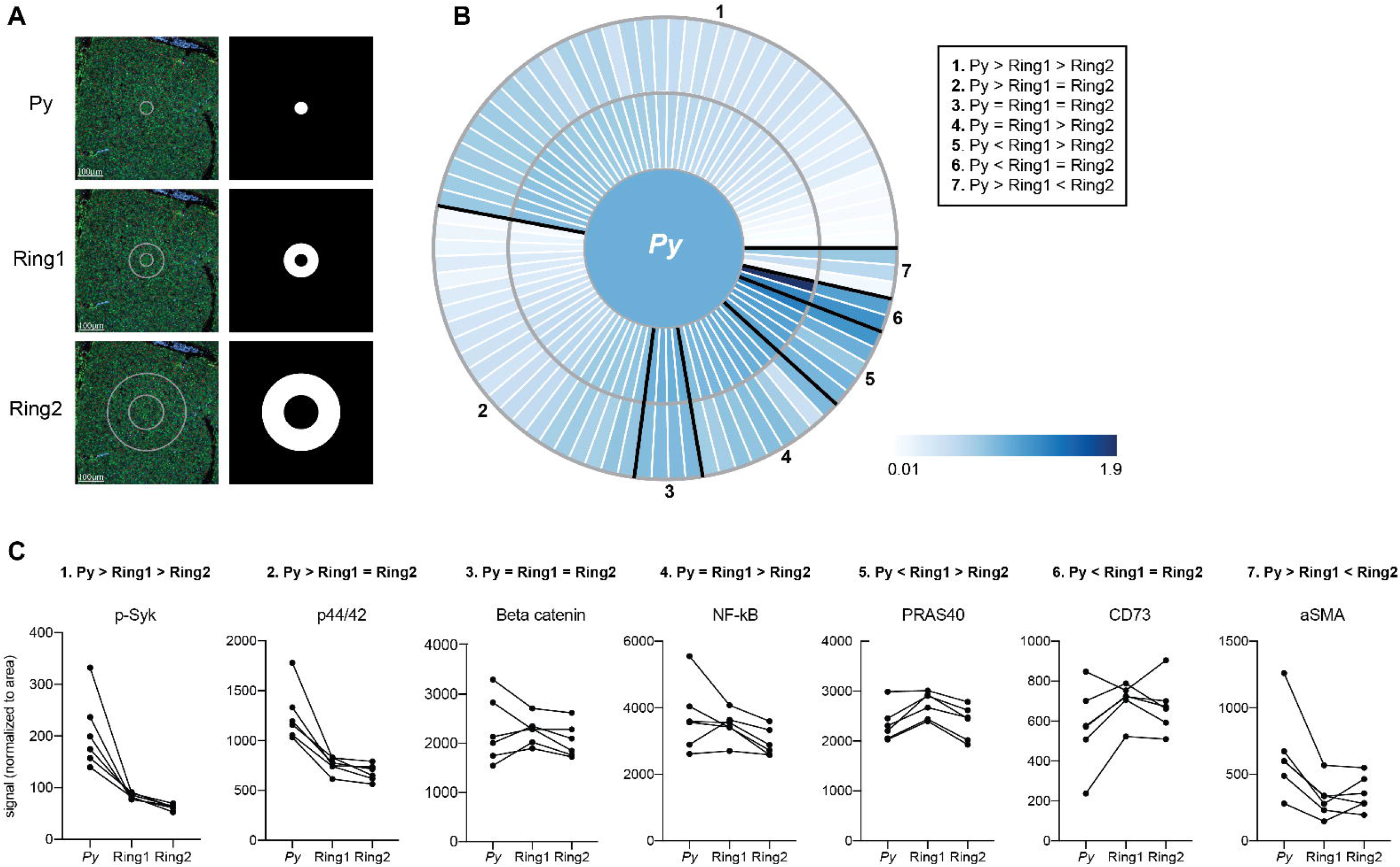
Signals are altered in rings surrounding *Plasmodium-infected* hepatocytes. (A) Representative images of fluorescent staining of liver sections and region of interest (ROI) masks. (B) Heat maps showing fold change in protein levels between in ring ROIs. Signal was normalized to ROI area and to the infected cell ROI, set at 1. Proteins showing similar relative patterns of expression across rings are grouped together and outlined in black. (C) Area normalized signal of two antibodies illustrative of delimited and gradual spatial patterns. Lines connect matched infected cell and ring ROIs. n= 6

Macrophages within the liver can be resident macrophages or monocyte-derived macrophages. Kupffer cells, the resident liver macrophage, are the most prevalent non-parenchymal cell in the liver, making up about 35% of total cells [38]. To investigate the distribution of Kupffer cells around infected hepatocytes, we stained liver sections with the Kupffer cell marker CLEC4F [37, 39, 40] and visualized liver stage parasites using DAPI (Fig 4A-C). We observed an increase in CLEC4F^+^ cells surrounding the parasite, with elevated density in Ring 1 compared to Ring 2, bystander cells within the same liver, and an identical area of tissue within uninfected animals (Fig. 4D). Interestingly, CLEC4F^+^ Kupffer cells often appear to wrap themselves around the LS-infected hepatocyte (Fig. 4B). We then asked if high Kupffer cell density around infected cells at 44 hpi could be due to selection of an existing microenvironment at the time of hepatocyte invasion, or due to cells migrating to the site after infection had been established. Although often referred to as “resident”, Kupffer cells have been shown to migrate along sinusoids within the liver (mean of 4.6μm/min) [41]. When we quantified Kupffer cells around *P. yoelii* parasites in livers collected 24 hpi, we observed no statistically significant difference between the number of cells in Ring1 and Ring2 (Fig 4E). Because parasites are much smaller at 24hpi than 44hpi, with average diameters of 10μm and 45μm respectively, we also measured Kupffer cell distance from the parasite membrane. The most notable difference in distribution between 24 and 44hpi was the increase in Kupffer cell density within 40μm of the parasite membrane (Fig 4F). To further explore the hypothesis that the parasite is surrounded by Kupffer cells that have migrated to Ring 1 between 24 and 44hpi, rather than a shifting of cells due to the increase in hepatocyte mass that occurs as a result of LS growth, we compared the average number of Kupffer cells within 5μm from the membrane of the 44h parasite and 22.5μm from the membrane of the 24h parasite (55μm diameter ROIs) (Fig S3). Despite the increased potential area that could be occupied by Kupffer cells within the 24h ROI due to the smaller volume occupied by the parasite, there were ~4.4x more Kupffer cells within the 44h ROI. This difference was maintained when the ROI diameter was expanded to 65μm (Fig S3), indicating that the increase in Kupffer cell density at 44hpi is not due to the expansion of the parasite towards pre-existing cells within close proximity.

**Figure 4.**
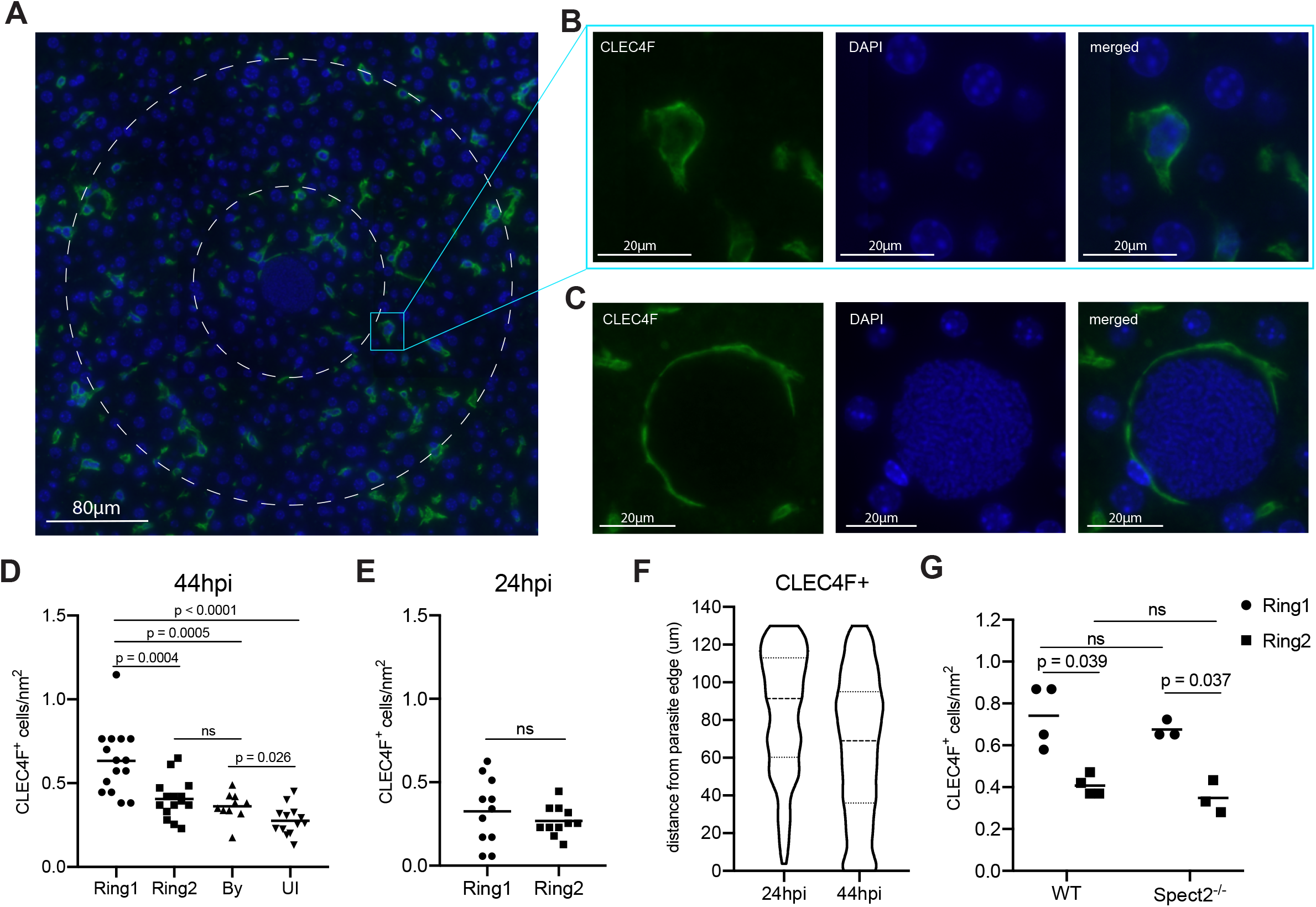
A high density of macrophages surrounds *Plasmodium* infected hepatocytes. (A) Representative image of CLEC4F^+^ staining within an infected liver. Nuclear DAPI staining is shown in blue, CLEC4F in green. Ring1 and Ring2 outer boundaries are indicated by dashed white circles. (B) CLEC4F staining of a single cell within Ring2 (C) Representative image of a CLEC4F^+^ cell in close proximity to a parasite at 44hpi. (D) Number of CLEC4F^+^ cells normalized by area in Ring1, Ring2, bystander (By) tissue, and uninfected (UI) tissue at 44hpi. 3-6 region of interest (ROI)s were counted from each of three mice. (E) Number of CLEC4F^+^ cells normalized by area in Ring1 and Ring2 at 24hpi. (F) Violin plot showing distribution of CLEC4F^+^ cells binned by distance from parasite edge at 24 and 44hpi. (F) Levels of CLEC4F^+^ cells in Ring1 and Ring2 around wild type (WT) and Spect2^-/-^ parasites at 44hpi.

Finally, to evaluate if a Kupffer cell dense region was selected as part of the sporozoite traversal process that occurs prior to hepatocyte entry, we utilized the spect2-parasite strain. Wild type parasites enter the liver through a hepatocyte, Kupffer cell, or liver endothelial cell, and then traverse through several hepatocytes using a transient vacuole before finally invading a final hepatocyte within a parasitophorous vacuole [3, 42, 43]. Spect2^-^ parasites that do not successfully invade are phagocytosed by Kupffer cells or fail to egress from their transient vacuole and are eliminated by host cell lysosomes [44–46]. This inability to traverse multiple hepatocytes limits their ability to travel through many cells in order to select a particular local microenvironment. Additionally, it limits the number of cells within the liver that come into direct contact with sporozoites. We infected mice with the spect2^-^ parasite strain and measured Kupffer cell density at 44hpi. The pattern of Kupffer cell density around infected hepatocytes was maintained in the context of spect2^-^ parasite infection (Fig 4F), indicating that cell traversal does not contribute to the Kupffer cell density around the infected cell.

We next sought to investigate the molecular characteristics of parasite-surrounding Kupffer cells. We revisited the DSP data (Fig. 3, Table S4), and calculated pairwise Pearson’s correlation coefficients between area normalized signal in Ring1 for all antibodies that were upregulated in Ring1 when compared to Ring2. We reasoned that if levels of two or more of these proteins correlated strongly with each other it could be because they are present within the same cells. Using Pearson’s correlation coefficients, we identified subsets of proteins that correlated with each other (Fig. S2). The strongest correlations were between B7H3, CD163, and Src, all of which are expressed by Kupffer cells and have been linked to tolerogenic M2 polarization of macrophages, particularly within the tumor microenvironment [35, 47–49].

We asked if the correlated proteins were expressed in overlapping populations of cells in Ring1 and Ring2. We found that CD163 was exclusively, and B7H3 almost exclusively, expressed on CLEC4F^+^ cells (Fig 5A-B). We also stained for PD-L1, which was part of both antibody panels and consistently upregulated in Ring1 compared to Ring2 but did not correlate with B7H3 and CD163. PD-L1 was found on both CLECF4^+^ and CLEC4F^-^ cells within Ring 1, with over 60% of PD-L1^+^ cells not expressing CLEC4F (Fig 5A-B). This is consistent with its lack of correlation with CD163 and B7H3, as well as published studies demonstrating PD-L1 expression on a variety of immune cell types [50]. B7H3^+^ and PD-L1^+^ Kupffer cells were very rare, 1.6% and 2.9% of all CLEC4F^+^ cells, respectively, however CD163^+^ cells were abundant and represented a majority of CLEC4F^+^ cells (Fig 5C).

**Figure 5.**
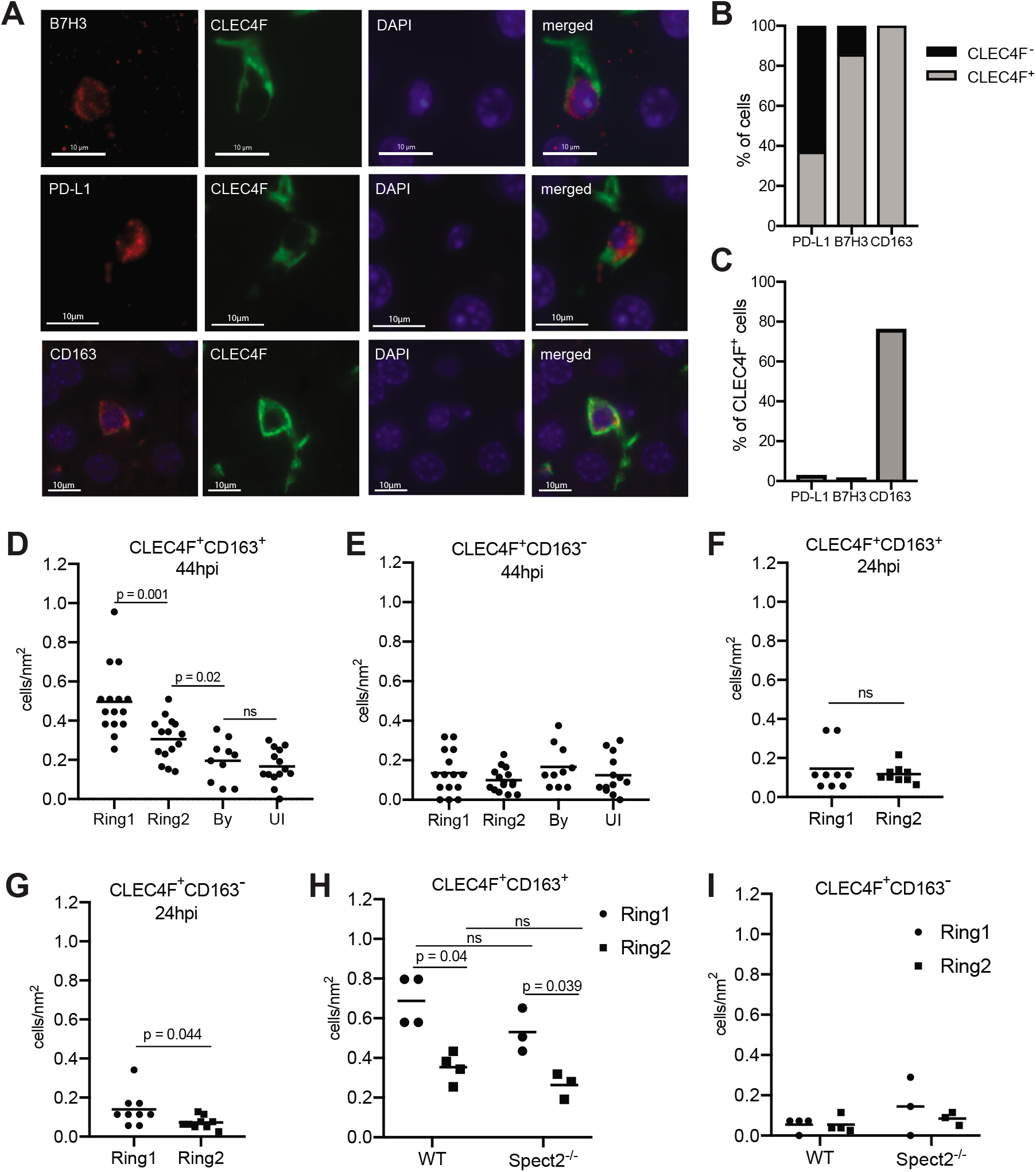
Tolerogenic macrophages migrate to surround *Plasmodium* infected hepatocytes. (A) Representative images of fluorescent staining of infected tissue for B7H3, PD-L1, and CD163. (B) Proportion of PD-L1^+^, B7H3^+^, and CD163^+^ cells within rings around infected cells that were CLEC4F^+^ or CLEC4F^-^. (C) Proportion of CLEC4F+ cells within rings around infected cells that were PD-L1^+^, B7H3^+^, or CD163^+^. (D) Number of CD163^+^CLEC4F^+^ and (E) CD163^-^CLEC4F^+^ cells normalized by area in Ring1, Ring2, bystander (By) tissue, and uninfected (UI) tissue at 44hpi. 3-6 regions of interest (ROIs) were counted from each of three mice. (F) Number of CD163^+^CLEC4F^+^ and (G) CD163^-^CLEC4F^+^ cells normalized by area at 24hpi. (H) Number of CD163^+^CLEC4F^+^ and (I) CD163^+^CLEC4F^+^ cells normalized by area, in Ring1 and Ring2 around wild type (WT) and Spect2^-/-^ parasites at 44hpi.

Quantification of CD163^+^ Kupffer cells revealed that more CD163^+^CLEC4F^+^, but not CD163^-^ CLEC4F^+^, cells were present in in Ring1 compared to Ring2, bystander, and uninfected tissue regions at 44hpi (Fig5D-E). We investigated the distribution of CD163^+^ Kupffer cells around *P. yoelii* parasites in livers collected 24 hpi and found no difference in CLEC4F^+^ CD163^+^ cell density between Ring1 and Ring2 (Fig 5F). Interestingly, at 24 h CLEC4F^+^ CD163^-^ cells were slightly elevated in Ring1 compared to Ring2 (Fig 5G). Finally, we compared CD163 expression in Ring1 and Ring2 around WT and Spec2-parasites. CLEC4F^+^ CD163^+^ cells were present at a higher density in Ring1 compared to Ring2 in both contexts, with no difference between parasite strains (Fig 5H). CLEC4F^+^ CD163^-^ cell levels were not significantly different between rings or between parasite strains (Fig5I).

## Discussion

In this study we utilized digital spatial profiling to characterize host total and phosphorylated proteins in and around *Plasmodium-infected* hepatocytes *in vivo.* Doing these analyses while preserving the tissue architecture allowed us to link a specific microenvironment to infected cells. Probing a large panel of proteins simultaneously in the same tissue regions allowed us to investigate the molecular characteristics of parasite-surrounding cells. Importantly, this is challenging to do with conventional approaches as it requires candidate-based investigation into specific candidate markers that may or may not be relevant for the cell type of interest. By measuring changes in host proteins in concentric rings around infected hepatocytes and correlations between these proteins we identified an influx of Kupffer cells towards the parasite and an increase in CD163 expression in these cells.

Kupffer cells originate from fetal liver erythromyeloid progenitors and in the adult liver, under resting conditions, their populations are self-renewing independent of bone marrow-derived cells [51]. Upon Kupffer cell depletion, infiltrating circulating monocytes differentiate into Kupffer cells starting 96 hours post-depletion [40]. Notably, monocyte-derived Kupffer cells do not begin expressing CLEC4F until between 72-96 hours post-depletion [40], indicating that the increase in CLEC4F^+^ cells we observed near the parasite between 24 and 44hpi cannot be due to differentiation of infiltrating monocytes and that it is almost certainly resident CLEC4F^+^ cells that have migrated towards the parasite.

Macrophages exist along a continuum of states that are often described as ranging from pro-inflammatory (M1) to tolerogenic (M2)[39]. Alterations of Kupffer cells upon sporozoite exposure have led to the hypothesis that parasites manipulate Kupffer cells to produce a tolerogenic environment for their development within the liver. Kupffer cell exposure to sporozoites has been shown to suppress respiratory burst [52], suppress antigen presentation [53], and skew cytokine production upon pro-inflammatory stimulation towards an anti-inflammatory response [54]. Co-culture of CD8^+^ T cells with sporozoite-stimulated monocyte-derived macrophages also produced less IFNγ [55]. Most of these studies were conducted with prolonged co-incubation of sporozoites and macrophages *in vitro.* Several functional studies have been conducted in which Kupffer cells are depleted before sporozoite infection [56, 57], but the importance of these cells for infection maintenance are confounded by the effects of depletion on hepatocyte invasion.

CD163 is commonly utilized as a tolerogenic macrophage marker [39], however it may also play a functional role in maintenance of LS infection. CD163 is a scavenger receptor expressed on monocytes and macrophages that binds and facilitates the internalization and clearance of hemoglobin-haptoglobin (HbHp) complexes, thereby protecting the liver from oxidative damage [58]. Binding of HbHp complexes promotes the expression of heme oxygenase-1 (HO-1) which degrades the Hb heme subunit, producing biliverdin, iron, and carbon dioxide. Although not part of our DSP antibody panel, HO-1 has been shown to be upregulated in macrophages and hepatocytes during *Plasmodium* LS infection and to be essential for infection maintenance [59]. Of particular interest, HO-1 was not found to be essential for *Plasmodium* LS infection when hepatocytes are cultured alone *ex vivo*, suggesting its effect on nonparenchymal cells influences infection. Higher expression of CD163 on Kupffer cells has also been linked to greater phagocytic activity [60]. Merozoite forms of the parasite exit the infected hepatocyte and enter the blood stream between 50-52 hpi. It is intriguing to speculate that the wrapping of Kupffer cells around infected hepatocytes (Fig 4C) could suggest a role for Kupffer cells in clean-up of the infected cell post-parasite exit. By regulating antigen presentation and inflammation around the infected cell microenvironment, Kupffer cells could be influencing the development of subsequent immunity. This could have long-reaching consequences not only for infection, but also for the development of whole parasite vaccines.

We are unable to determine from our data if CD163^+^ Kupffer cells are infiltrating in towards the parasite, or if they begin expressing CD163 upon gaining their location near the infected cell, however the small increase in CD163^-^CLEC4F^+^ cells in Ring1 compared to Ring2 at 24 hpi (Fig. 5G) supports the latter hypothesis. PD-L1 expression, which is increased in Ring1 compared to Ring2, has been shown to be induced in monocyte-derived macrophages in the skin upon exposure to *Plasmodium* sporozoites [55]. A portion of sporozoites are thought to cross and interact with Kupffer cells as they are entering the liver [43], however, as no difference in CD163^+^ Kupffer cell density was observed between WT parasites and Spec2^-^ parasites, which are traversal deficient, we hypothesize that the increase in CD163 expression is unlikely to be triggered by pre-invasion events.

By utilizing DSP we were able to measure host protein on a large scale in single infected hepatocytes *in vivo* without sorting cells and without dissociating cells from their microenvironment. While we cannot rule out non-specific binding of antibodies to parasite proteins, the minimal sample processing may better preserve the host cell condition compared to experiments that require hours of cell sorting, particularly in the case of post-translational modifications. Several of the proteins up-regulated in infected cells were phosphorylated, indicating increased activity: p-IK-Ba, p-S6, and p-Erk. Consistent with our results, Erk (MAPK1) activity was previously identified by our lab as important for maintenance of *P. yoelii* infection *in vitro* using a kinase inhibitor screen combined with a machine learning algorithm [23]. Additionally, levels of p-S6 are higher in hepatocytes that are more susceptible to *Plasmodium* infection and in infected hepatocytes *in vitro*, although in the context of infection S6 phosphorylation is dysregulated from classical upstream activator p-Akt [11]. One surprising result was the increase seen in p53 levels in infected cells. P53 is suppressed in infected hepatocytes and this suppression is essential for maintenance of infection *in vitro* and *in vivo*, however, in these studies p53 levels were not measured past 24hpi [12]. We hypothesize that the increase in p53 seen here at 44hpi could be indicative of a loss of regulation by the parasite as it shifts towards merozoite production and preparation for egress.

A very small number of parasites successfully invade hepatocytes and complete LS infection. This, and the extensive remodeling of infected hepatocytes, suggest *Plasmodium* parasites have substantial requirements of their host cells. By identifying proteins/post-translational modifications that show very little variation among infected compared to uninfected cells, we may be able to identify specific targets or signaling nodes that are maintained within, or selected for, very narrow limits by the parasite. These factors could represent promising drug targets, as even small perturbations of these factors could have dire consequences for the developing LS parasite. While this work is entirely focused on the rodent parasite *Plasmodium yoelii*, DSP is readily adaptable to the study of human-infectious species *Plasmodium falciparum* and *Plasmodium vivax* in the recently developed humanized mouse model [61, 62]. Evaluating the spatially resolved host transcriptomic and proteomic responses that occur after infection, particularly in the context of the dormant *P. vivax* hypnozoite, may reveal novel biology regulating infection maintenance and development of immunity.

## Methods

### Mosquito rearing and sporozoite production

Female 6–8-week-old Swiss Webster mice (Harlan) were injected with blood stage *Plasmodium yoelii* 17XNL parasites. Infected mice were used to feed female *Anopheles stephensi* mosquitoes after gametocyte exflagellation was observed. Salivary gland sporozoites were isolated according to the standard procedures at days 14 or 15 post blood meal. Animal handling was conducted according to the Institutional Animal Care and Use Committee-approved protocols.

### Mouse infections

6-8-week-old female Balb/cAnN mice were purchased from Envigo. All mice were maintained in accordance with protocols approved by Seattle Children’s Research Institute Institutional Animal Care and Use Committee (IACUC). Mice were infected by retro orbital injection with 100,000 or 1 million *P. yoelii* sporozoites. Livers from infected, or uninfected age-matched, mice were harvested at 24 or 44 hpi and fixed in 4% paraformaldehyde for 24 hours. Tissues were then paraffin embedded, cut into 4mm sections, and mounted on positively charged glass slides. Mounted liver slices were then used for digital spatial profiling or immunofluorescence staining.

### Digital spatial profiling

Digital spatial profiling (DSP) was performed by NanoString Technologies using the GeoMx Digital Spatial profiler. For selecting regions of interest, slides were stained with DAPI and a fluorescent conjugated antibody against *Py*HSP70. Slides were simultaneously incubated with one of two pre-validated panels of 42-43 oligo-tagged antibodies (Table S1). Counts were normalized to an internal control (ERCC) and to ROI area. Data were analyzed by ANOVA with paired or unpaired multiple comparisons as appropriate. Our single infected-cell ROIs may encompass a portion of neighboring cells.

### Immunofluorescence staining

Slide-mounted liver slices were washed twice in xylene for 3 minutes followed by washes in 100%, 95%, 70% and 50% ethanol for 3 minutes each. Slides were then washed with DI water and heated to 90C for 30 minutes in 1% citrate-based antigen unmasking solution (Vector Laboratories) using a Biocare Medical Decloaking Chamber. Slides were washed with TBS-0.025% Tween (TBST) and then blocked for 4 hours in TBST containing 1.5% BSA and 15% goat serum (Sigma Aldrich). Slides were incubated in primary antibodies at 4C overnight. Following primary antibody staining, slides were washed with TBST and incubated with secondary antibodies and DAPI (1:3,000) for 1 hour at room temperature. Slides were washed with TBST and autofluorescence quenched using Vector TrueView (Vector Labs). Fluoromount G mounting media was used to preserve fluorescence signal. Primary antibodies were used at the following concentrations: *Py*Hsp70 1:1,000, *Py*CSP-488 1:500, p-p44/42 1:200 (Cell Signaling 4370), p-IK-Ba 1:200 (Cell Signaling 2850), p-Akt 1:100 (Cell Signaling 9271), CD163 1:500 (Proteintech 16646-1-AP), CLEC4F-647 1:100 (BioLegend 156804), PD-L1 1:200 (Cell Signaling 64988), B7H3 1:200 (Novus Bio NB600-1441). Secondary antibodies anti-mouse AlexaFluor-488, anti-rabbit AlexaFluor-594, and anti-rabbit AlexaFluor-647 (Invitrogen) were used at a 1:1,000 dilution.

### Imaging and quantification

Images (40X) were acquired using a DeltaVision Elite High Resolution Microscope. Z-stacks of 0.3*μ*m thickness were taken for images encompassing infected and uninfected cells. For cell quantification within Ring ROIs 3×3 image panels were taken with a 60-pixel overlap. Images were stitched and deconvolved using the DeltaVision Softworx software and were visualized using Imaris software. ImageJ was used to quantify fluorescence intensity within defined ROIs. Distances from parasites to Kupffer cells were measured between nucleus centers, or from the parasite membrane to the Kupffer cell nucleus, using Imaris software. Only Kupffer cells with a visible, stained nucleus (DAPI) were included in counts.

The phosphosignaling network was reconstructed using PhosphoSitePlus®, a curated knowledgebase dedicated to mammalian post-translational modifications (https://www.phosphosite.org) [63].

## Acknowledgments

We would like to acknowledge Liuliu Pan and Yan Liang at NanoString Technologies for their assistance with our DSP runs. This work was funded by R01GM101183 from the National Institutes of Health to AK. SR is the recipient of T32 training grant 5T32HD007233-39 from the University of Washington.

---

**Fig S1**. **DSP results are reproducible across runs and between pooled and single ROIs.** Average fold change between infected and uninfected ROIs for each antibody from (A) two independent DSP runs, and (B) two slides run at the same time. Data were analyzed by linear regression. (C) For each antibody the average area-normalized signal 9 single infected ROIs was plotted against that of one pooled infected ROI. (D) For each antibody the average area-normalized signal 9 single uninfected ROIs was plotted against that of one pooled uninfected ROI. Data were analyzed by linear regression.

**Fig S2**. **A subset of upregulated (phospho)proteins in proximity to *Plasmodium-infected* hepatocytes are correlated.** (A) Heat map indicating the Pearson correlation coefficient for each pair of antibodies for those significantly upregulated in Ring1 compared to Ring2. n = 6.

**Fig S3. Growth of *Plasmodium* infected hepatocyte does not account for increased Kupffer cell density around cell.** Kupffer cell density within circular ROIs of 55um and 65um around parasites at 24hpi and 44hpi. Parasites are shown as green circles. Length of lines is indicated in microns. Circles and rings are shown to scale. Kupffer cell density is shown as the mean from 3-4 parasites per mouse from 3 mice per time point.

