## Supplementary figures and images for "Elucidating spatially-resolved changes in host signaling during *Plasmodium* liver-stage infection"

### Figure S1

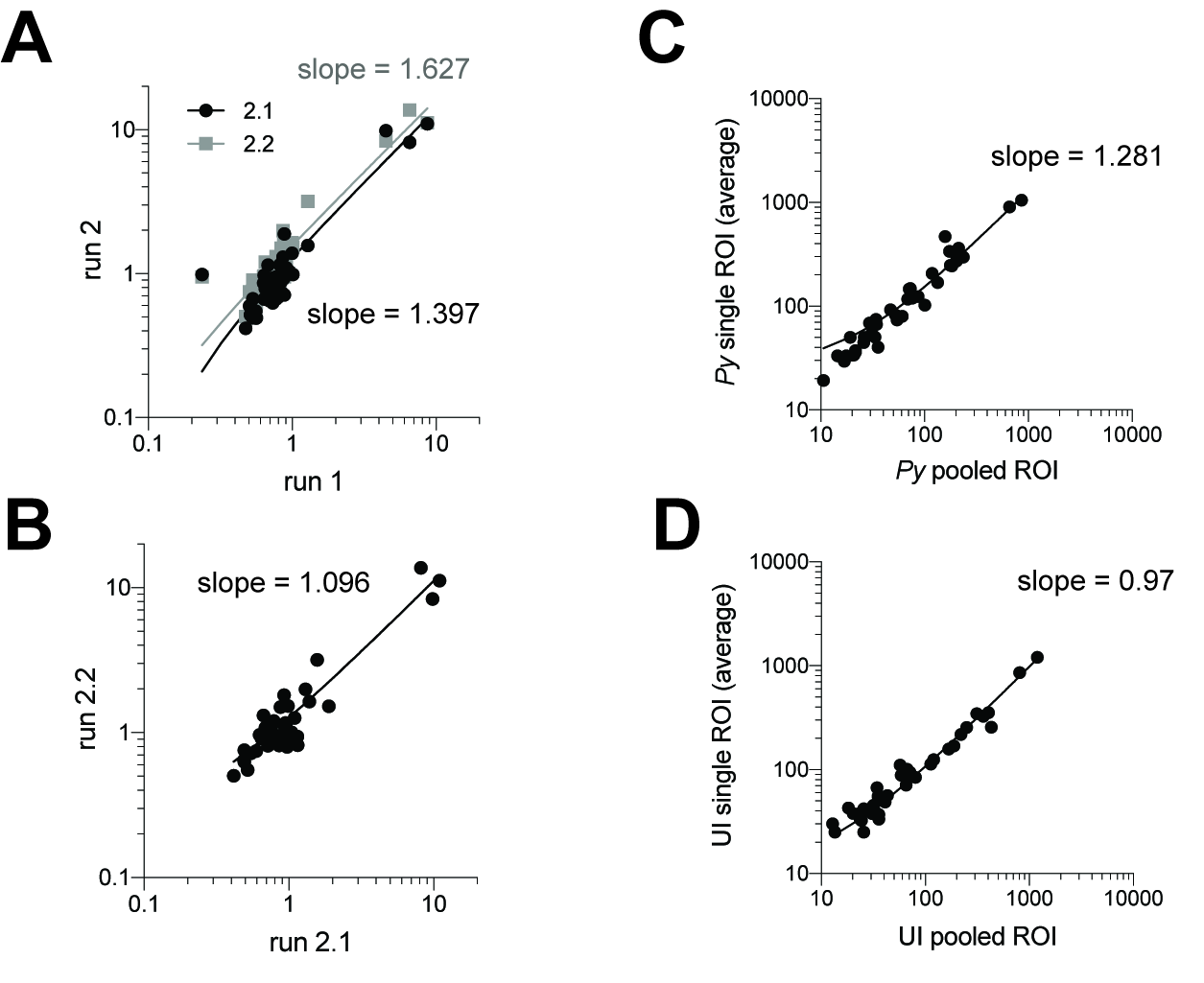

### Figure S2

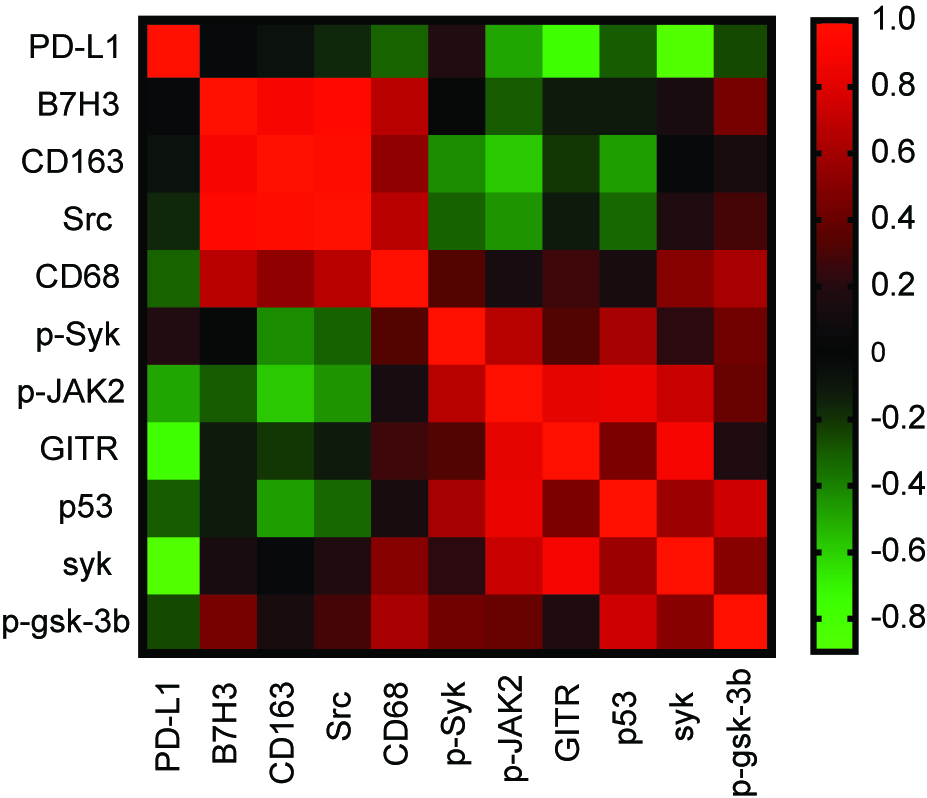

### Figure S3

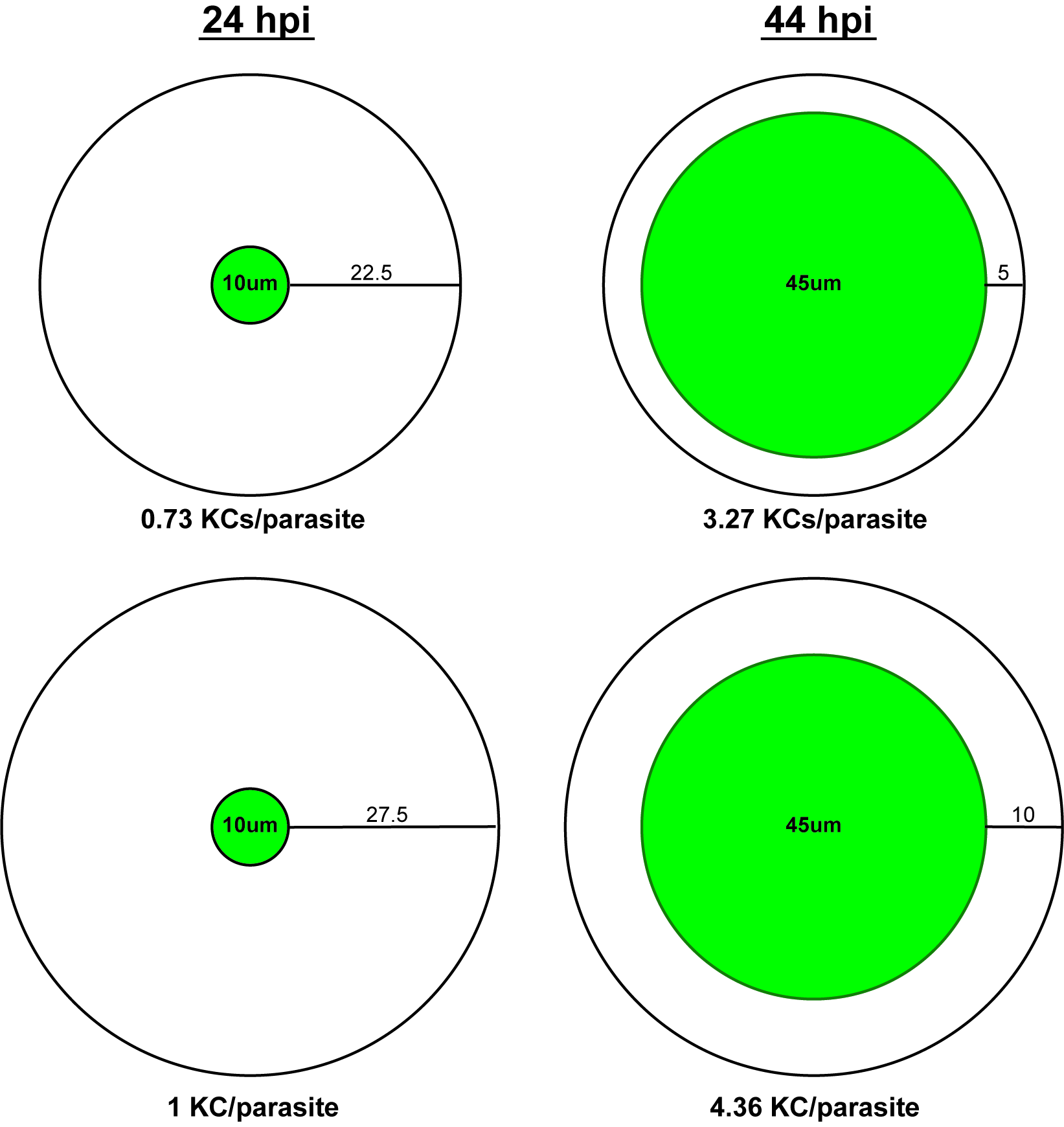
